# Natural Language Processing Applied to Spontaneous Recall of Famous Faces Reveals Memory Dysfunction in Temporal Lobe Epilepsy Patients

**DOI:** 10.1101/2024.08.23.609193

**Authors:** Eden Tefera, Helen Borges Delfino de Souza, Charlotte Blewitt, Aaqib Mansoor, Haley Peters, Peem Teerawanichpol, Simon Henin, William B. Barr, Stephen B. Johnson, Anli Liu

## Abstract

**Objective and Background:** Epilepsy patients rank memory problems as their most significant cognitive comorbidity. Current clinical assessments are laborious to administer and score and may not always detect subtle memory decline. The Famous Faces Task (FF) has robustly demonstrated that left temporal lobe epilepsy (LTLE) patients remember fewer names and biographical details compared to right TLE (RTLE) patients and healthy controls (HCs). We adapted the FF task to capture subjects’ entire spontaneous spoken recall, then scored responses using manual and natural language processing (NLP) methods. We expected to replicate previous group level differences using spontaneous speech and semi-automated analysis.

**Methods:** Seventy-three (N=73) adults (28 LTLE, 18 RTLE, and 27 HCs) were included in a case-control prospective study design. Twenty FF in politics, sports, and entertainment (active 2008-2017) were shown to subjects, who were asked if they could recognize and spontaneously recall as much biographical detail as possible. We created human-generated and automatically-generated keyword dictionaries for each celebrity, based on a randomly selected training set of half of the HC transcripts. To control for speech output, we measured the speech duration, total word count and content word count for the FF task and a Cookie Theft Control Task (CTT), in which subjects were merely asked to describe a visual scene. Subjects’ responses to FF and CTT tasks were recorded, transcribed, and analyzed in a blinded manner with a combination of manual and automated NLP approaches.

**Results:** Famous face recognition accuracy was similar between groups. LTLE patients recalled fewer biographical details compared to HCs and RTLEs using both the gold-standard human-generated dictionary (24%±12% vs. 31%±12% and 30%±12%, p=0.007) and the automated dictionary (24%±12% vs. 31%±12% and 32%±13%, p=0.007). There were no group level differences in speech duration, total word count, or content word count for either the FF and CTT to explain difference in recall performance. There was a positive, statistically significant relationship between MOCA score and FF recall performance as scored by the human-generated (ρ= .327, p= .029) and automatically-generated dictionaries (ρ= .422, p= .004) for TLE subjects, but not HCs, an effect that was driven by LTLE subjects.

**Discussion:** LTLE patients remember fewer details of famous people than HCs or RTLE patients, as discovered by NLP analysis of spontaneous recall. Decreased biographical memory was not due to decreased speech output and correlated with lower MOCA scores. NLP analysis of spontaneous recall can detect memory dysfunction in clinical populations in a semi-automated, objective, and sensitive manner.

## INTRODUCTION

Epilepsy patients rank memory problems as their most significant cognitive comorbidity, impacting daily function and school and workplace participation ^1^. Despite rapid gains in the fields of cognitive and computational neuroscience, clinical neuropsychological testing has remained largely unchanged ^2^. The advantage of standardized testing is its validation on large populations and normalized performance scores by age and education, However, the test administration and scoring process is laborious and yields an oversimplified measure of behavior. The development of novel, precise, and clinically meaningful approaches is needed for early and serial memory assessment in epilepsy, Alzheimer’s Disease ^2^, and other memory impaired patient populations.

Memory deficits are commonly observed in patients with Temporal Lobe Epilepsy (TLE) but are inconsistently captured by standard clinical testing ^3^. Due to their extensive connections with widespread cortical regions, the hippocampus and connected limbic regions, are hijacked by seizure networks ^4^. The Rey Auditory Verbal Learning Test (RAVLT), created in 1941, is the most widely used test of verbal memory function in assessment of TLE patients ^5^. Poorer RAVLT performance for left TLE patients, as compared to right TLE patients, has been well established ^6,7^. While the test is a useful predictor of seizure laterality, it may be insensitive to subtle impairment over time and performance is influenced by executive and language ability.

Cognitive testing in clinical settings could embrace more naturalistic behaviors and utilize computational methods to measure memory deficits more efficiently and objectively. Cognitive neuroscience has already embraced more realistic behavioral paradigms, such as spontaneous speech ^8^, autobiographical recall ^9^, film watching ^10^, and physical navigation ^11,12^. Similarly, computational methods including Natural Language Processing (NLP) have been used to quantify distinct speech components, including lexicon and syntax, to distinguish patients with Alzheimer’s Disease and Mild Cognitive Impairment ^13–17^ from healthy controls and to predict progress to psychosis among at-risk youth.

We adapted the Famous Faces (FF) Task to capture and analyze the spontaneous recall from TLE patients and healthy controls, in a case-controlled prospective study design. The Famous Faces task was designed in the 1970s to assess face recognition and biographical memory for a set of public figures. While initially developed for assessment of amnestic patients with Korsakoff’s syndrome, the task has consistently shown that patients with LTLE demonstrate poorer remote recall for famous names and biographical details compared to healthy controls and patients with RTLE ^18–21^. Previously, RTLE patients have been demonstrated to have poorer facial recognition ^22^, although performance on non-verbal memory tasks has been variable ^23^. Our goal was to measure memory performance by analyzing subjects’ spontaneous recall through both human and semi-automated approaches using NLP. We hypothesized that patients with LTLE, whose seizures likely affect mesial temporal regions involved in episodic and semantic memory, would spontaneously recall fewer details than healthy controls and RTLE patients and that this would be distinct from differences in speech output.

## METHODS

This study was conducted following protocols approved by the New York University Institutional Review Board. All study activities complied with regulations for human subject research, and all data was collected during a single study session.

### Eligibility criteria

We recruited Temporal Lobe Epilepsy (TLE) subjects and Healthy Controls (HCs) ages 18-60 from a single Level 4 Epilepsy Center from 2018-2023. HCs were included if they were between the ages of 18 and 60, did not have a self-reported history of neurological or psychiatric disease, and earned a normal score on the Montreal Cognitive Assessment (MOCA >=26/30 ^24^). The MOCA is a widely used cognitive screening tool assessing multiple cognitive domains, including memory, attention, executive function, visuo-spatial construction, naming and orientation ^25^. Patients with temporal lobe epilepsy who scored >= 22/30 on the Montreal Cognitive Assessment were included. A lower threshold for TLE patients was chosen to include patients with objective memory impairment and to assess variability in recall performance in our famous faces task. Epilepsy localization was determined by seizure semiology, MRI Brain, and EEG concordance, and adjudicated by a board-certified neurologist and epileptologist. Only patients with a probable or definite focal epilepsy localized to unilateral temporal lobe were included (meaning at least two concordant criteria without discordant criteria).

### Sample size estimates

Sample size estimates were based on previously published result demonstrating that patients with LTLE have poorer naming of familiar celebrity faces compared to healthy controls ^26^. For two independent study groups (assuming HCs and LTLE as primary comparison) with a continuous endpoint (percentage of detailed recalled of recognized celebrities), we calculated that a sample size of 17 subjects per group would be adequate to detect a large effect size (power 90%, alpha 0.05).

### Famous Faces Task and Cookie Theft Control Task

The Famous Face Test was adapted from the Iowa Famous Face Test ^18^. The test is designed to assess remote memory for face naming and face recognition abilities. **(Fig 1)**. The test includes two phases: (1) the familiarity, naming, and spoken recall of 20 famous faces **(Fig 1a**), and (2) the recognition of famous faces in a multiple-choice format **(Fig 1b)**, similar to prior famous face studies ^21,26^. All subjects were exposed to the same 20 celebrity faces in the same order and shown the same multiple-choice tests.

**Fig 1.**
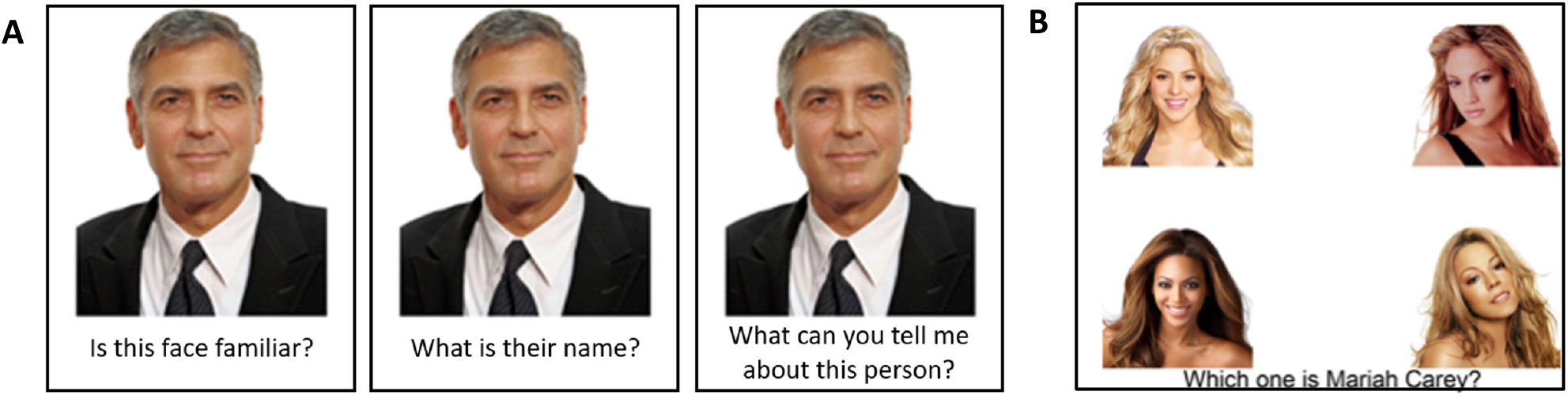
Famous Face Task.

To create the set of celebrities, we used the MIT Media Lab’s Pantheon Dataset of Historical Popularity that ranked famous individuals by year ^27^. We first selected 98 famous individuals from entertainment, politics, sports, and music who were born in the U.S. between 1960s-2000s and were well-known in the decade prior to the initiation of the study (2008-2017). An online Qualtrics questionnaire containing these names was sent out to 44 healthy participants (ages 18-50) with the question, “Which of these famous individuals can you identify based on their photos?” We excluded famous individuals that were recognized by less than 60% of healthy participants. We then selected a list of 45 individuals to be used in the Famous Face Test, with 20 celebrities used for free recall, and the remainder of faces were used in the multiple--choice component of the task.

To address the potential bias of uneven exposure to popular culture, only celebrities that were recognized by each subject were included in analysis. To control for potential speech and language impairment in TLE patients ^28^, we added the Cookie Theft Task from the Boston Diagnostic Aphasia Examination (BDAE) ^29^ after the study was initiated. Subjects’ responses were recorded and stored on the local HIPAA-compliant servers. Subjects were tested in one of two settings: (1) on-site at the NYU Comprehensive Epilepsy Center or (2) remotely via WebEx, a HIPAA compliant desktop conference call application. Webex was added as a testing platform during the COVID-19 pandemic when all research was conducted remotely.

### Speech Transcription

Subjects spontaneous recall responses to each of the 20 celebrity faces were recorded and transcribed in two ways: (1) human transcription and (2) WebEx transcripts generated after a recording session with manual review. For subjects participating on-site, WebEx transcriptions were retroactively generated using the original audio files. WebEx automatically transcribes audio of meetings recorded in the MP4 format. The WebEx-generated transcripts included time stamps and were verified for accuracy by a human reviewer. Subject and interviewer speech were manually separated by the human rater to ensure that transcripts only contained transcribed speech from the subject.

### Speech Analysis

Subjects’ transcripts were analyzed using computer code written in the Python language using spaCy, an open-source library for Natural Language Processing ^30^. The spaCy library takes unstructured text as input and returns structured output with extensive linguistic information. In particular, the library divides the text into tokens, which consist of words, numbers, punctuation and other symbols, and identifies the part of speech. Total word count and content word count were obtained for both sets of transcripts collected from FF and CTT, where total word count is defined as the number of tokens in a piece of text, and content word count is defined as the number of words containing the following parts of speech: noun, verb, adjective or adverb.

### Creation of Human-Generated and Automated Keyword Dictionaries and Subject Scoring

Two unique keyword dictionaries (human-generated and automated) were created for each celebrity. To avoid overfitting, we randomly selected half of the sample of the transcripts of the healthy controls (N=14). The human-generated keyword dictionary was created by two independent raters (ET and AM) who extracted key biographical details about each celebrity from the transcripts. The two human dictionaries were merged by including: (1) keywords present on both dictionaries included (2) keywords on either dictionary mentioned by two or more subjects and (3) keywords of similar meanings found on both lists (simplest derivative listed *ex: pass listed to represent passed away & passing)*.

The automated keyword dictionary was generated by pooling the randomly selected half of the HC transcripts for each famous person, creating 20 documents. Potential keywords for each celebrity were scored using Term Frequency-Inverse Document Frequency (TF-IDF), which measures the importance of a term within a document relative to the collection of documents ^31^. Word sequences (n-grams) were generated from the documents and filtered using orthography and part of speech. N-grams up to length 5 were selected when words were capitalized, and up to length 2 for lowercase words. Both sets required the presence of content words. The n-grams were scored using term frequency (the number of occurrences of a term within the document about a particular famous person) and inverse document frequency (the reciprocal of the number of famous people that share the term). The top 10% of the highest scoring n-grams were selected. When terms overlapped, the longer term was retained. *ex: if “George” and “George Clooney” were identified as potential terms, only the latter was kept*. The algorithm selected 3-13 keywords for each famous person, with an average of 8. Examples of human generated and automated keyword dictionaries are shown in **Table S1**.

Subjects were scored in by two reviewers on the percentage of keywords recalled for each recognized celebrity from both the human generated dictionary (gold standard) and automated dictionary. Scorers were blinded to the subject diagnosis and adjudicated when there was disagreement.

### Neuropsychological Testing

To screen for initial eligibility, all subjects were administered the Montreal Cognitive Assessment (MOCA) ^25^. Scores from a comprehensive neuropsychological test battery were available for a subset of TLE patients undergoing pre-surgical evaluation (n=18). Full Scale IQ was evaluated through the Test of Premorbid Functioning (TOPF) and the Verbal Comprehension Index (VCI) from the Wechsler Adult Intelligence Scale (WAIS-IV) ^32,33^. Verbal memory was evaluated through the Rey Auditory Verbal Learning Test (RAVLT) long delayed free recall score ^5^.

## Statistical Analysis

We performed descriptive statistics on the demographics and neuropsychological metrics for the 3 subject groups (LTLE, RTLE, HC), by calculating means and standard deviations for continuous measures (age, MOCA, TOPF, IQ, and RVLT) and counts for categorical measures (sex, handedness, and educational level). The Shapiro-Wilk test was used to test for normality of distribution for continuous data. Group level differences were calculated by the Kruskal-Wallis tests for continuous data and chi-square for categorical data. Descriptive statistics were calculated separately for subjects participating in the Famous Face Task (LTLE 28, RTLE 18, HC 27) and the subgroup of subjects who completed the control Cookie Theft Task (LTLE 17, RTLE 13, HC 23).

For famous face results, means and SDs were calculated for all continuous data, including famous face recognition, recalled biographical details from human dictionary recalled biographical details from automated dictionary, total word count, content word count, and speech duration. Only FF identified as familiar by subjects were included to obtain a keyword performance score. The primary outcome for this study was the percentage of details recalled for selected 20 celebrities as scored by the human-generated keyword dictionary. Percentage recalled was calculated for each subject, then averaged across diagnostic category (HC, LTLE, RTLE). Secondary outcomes included percentage of details recalled by group as scored by the automated keyword dictionary. In control analyses, speech output was measured by the total number of spoken words, content words, and speech duration during the FF and Cookie Theft Task. For all continuous data, we assessed the distribution of the data with the Shapiro-Wilk test and performed descriptive statistics (mean, SD). To compare group differences in recall performance between 3 independent groups, we used the Kruskal-Wallis tests, then the Wilcoxon rank-sum for post-hoc pairwise comparisons. We used a Spearman’s correlation test to compare remote biographical memory as measured by our keyword dictionaries and measures from validated neuropsychological tests (including MOCA and RVLT scores). Post-hoc effect sizes were calculated based on the primary and secondary outcome and reported as Cohen’s d.

## RESULTS

### Subjects

Seventy-three (73) adults completed the Famous Face Task: 28 LTLE, 18 RTLE, and 27 HC **(Table 1)**. There were no group-level differences in sex (60% F), handedness (85% RH), education status (70% college or above). Compared to TLE patients, HCs were younger (p= .018) and scored slightly better on the MOCA (p=.0001), which may be an artifact of the higher MOCA cutoff scores for HC eligibility. Within TLE patients, there were no group-level differences between LTLE and RTLE patients in MOCA, WAIS-IV FSIQ, or TOPF scores. However, LTLE patients had poorer performance on the RAVLT than RTLE patients (8.36±3.0 vs 11.50 vs 2.51, p=.023). All subjects spoke for an average of 766.7 seconds (SD 502.05 seconds). Fifty-three of the subjects who completed the FF task also completed the Cookie Theft Task (17 LTLE, 13 RTLE, and 23 HC) **(Table 2)**. There were no group-level differences in sex (68% F), handedness (85% RH), or education status in this subset of subjects. Compared to TLE patients, HCs were younger (p= .037) and had higher MOCA scores (p=.0001). There were no differences in MOCA scores between LTLE and RTLE patients.

**Table 1.** Famous Face Demographics (n=73)

|  |  | LTLE (n = 28) | RTLE (n = 18) | Healthy (n = 27) | Total (n = 73) | P value |
| --- | --- | --- | --- | --- | --- | --- |
| Age | Mean | 31.14 | 33.94 | 28.20 | 30.82 | *0.02 |
|  | (± SD) | 9.12 | 7.30 | 9.00 | 8.83 |  |
| Sex | Female | 17 | 11 | 18 | 46 | 0.88 |
|  | Male | 11 | 7 | 9 | 27 |  |
| Handedness | Right | 23 | 14 | 25 | 62 | 0.32 |
|  | Left | 4 | 3 | 1 | 8 |  |
|  | Ambidextrous | 1 | 1 | 0 | 2 |  |
| Education | ≤ 12 years | 4 | 0 | 0 | 4 | 0.24 |
|  | 13 - 15 years | 5 | 3 | 5 | 13 |  |
|  | 16 years | 11 | 7 | 9 | 27 |  |
|  | ≥ 17 years | 8 | 6 | 10 | 24 |  |
| Montreal Cognitive Assessment | Mean | 26.50 | 26.80 | 28.77 | 27.33 | *P<.001 |
|  | (± SD) | 2.32 | 2.32 | 1.14 | 2.28 |  |
| Test of Pre-morbid Functioning | Mean | 98.00 | 104.50 | N/A | 100.26 | 0.85 |
|  | (± SD) | 24.77 | 10.45 | N/A | 20.86 |  |
| IQ | Mean | 100.44 | 102.64 | N/A | 101.33 | 0.07 |
|  | (± SD) | 19.18 | 15.68 | N/A | 17.55 |  |
| Rey Auditory Verbal Learning | Mean | 8.36 | 11.50 | N/A | 9.50 | *0.02 |
|  | (± SD) | 3.00 | 2.51 | N/A | 3.17 |  |
| * = Significant Results |  |  |  |  |  |  |
| N/A= Not Assessed |  |  |  |  |  |  |

**Table 2.** Cookie Theft Demographics (n=53)

|  |  | LTLE (n = 17) | RTLE (n = 13) | Healthy (n = 23) | Total (n = 53) | P value |
| --- | --- | --- | --- | --- | --- | --- |
| Age | Mean | 30.30 | 33.31 | 27.62 | 29.96 | *0.03 |
|  | (± SD) | 7.41 | 7.15 | 8.60 | 8.04 |  |
| Sex | Female | 12 | 9 | 15 | 36 | 0.96 |
|  | Male | 5 | 4 | 8 | 17 |  |
| Handedness | Right | 14 | 10 | 21 | 45 | 0.28 |
|  | Left | 3 | 2 | 1 | 6 |  |
|  | Ambidextrous | 0 | 1 | 0 | 1 |  |
| Education | ≤ 12 years | 1 | 0 | 0 | 1 | 0.73 |
|  | 13 - 15 years | 3 | 2 | 4 | 9 |  |
|  | 16 years | 7 | 3 | 7 | 17 |  |
|  | ≥ 17 years | 6 | 6 | 9 | 21 |  |
| Montreal Cognitive Assessment | Mean | 26.69 | 26.13 | 28.91 | 81.73 | *P<.001 |
|  | (± SD) | 2.39 | 2.13 | 1.04 | 5.56 |  |

### LTLE subjects recalled fewer details for familiar FF compared to HCs and RTLE subjects

There were no group-level differences in FF recognition in the forced choice recognition portion of the FF task ((χ^2^ (2, N = 73) = 1.98, p = .780) **(Table 2, Fig S2)**, suggesting that exposure to famous faces could not account for differences in recall performance across groups. Recall performance differed between groups when scored against the human generated keyword dictionary (χ^2^ (2, N = 73) = 9.94, p = .007, **Table 2)**. Post-hoc pairwise comparisons showed that LTLE subjects recall fewer human-generated keywords than HCs (24±12% vs. 31±12%, d =0.58, p=.003) and RTLE subjects (30±10%, p= .005) for familiar FF **(Fig 3a**). Group-level differences in memory performance were also observed when scored by automatically generated keywords, (χ^2^ (2, N = 73) = 9.850, p = .007, **Table 2**). Post-hoc pairwise comparisons showed that LTLE subjects recall fewer automatically-generated keywords than HCs (24±12% vs 32±13%, d=0.64, p=.002, **Fig 3b)**.

**Fig 2.**
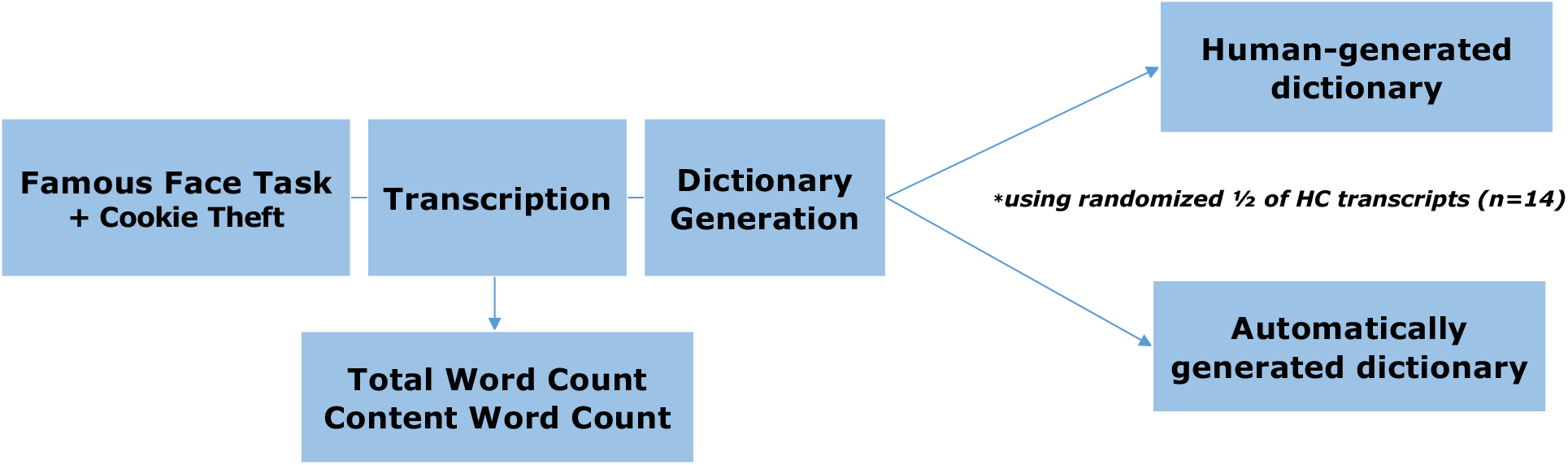
Dictionary Generation.

**Fig 3.**
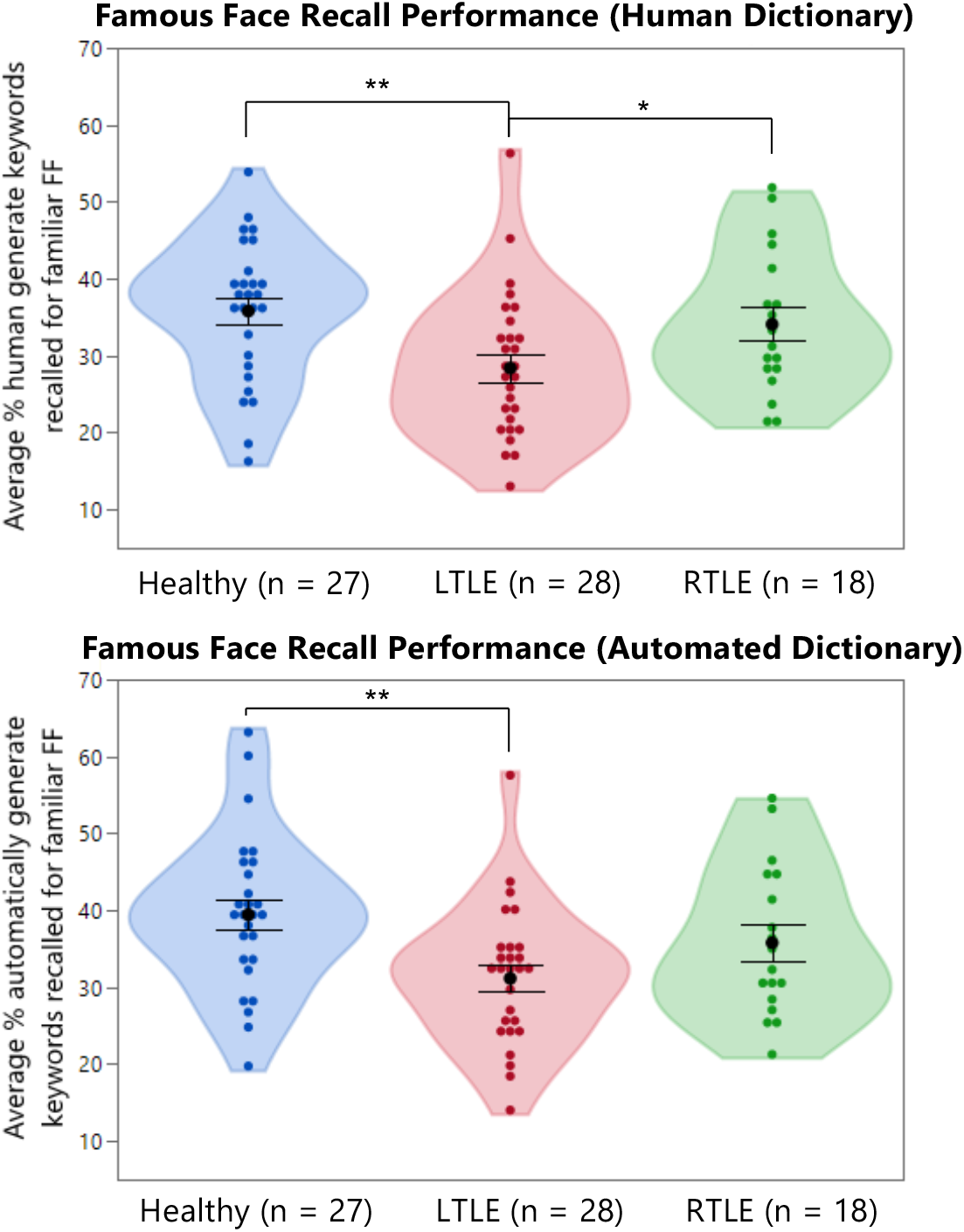
Famous Face Recall Performance. 3a. Human-generated Dictionary 3b. Automated Dictionary

### No group-level differences in speech output or FF exposure

There were no group level differences in speech duration for the Famous Face task (p=.175, **Table 2**) or the Cookie Theft task (p=.8063, **Table 3)**. For the Famous Face task, there were no group level differences in total word count (χ^2^ (2, N = 73) = 1.98, p = .372) or content word count (χ^2^ (2, N = 73) = 2.16, p = .340, **Table 2)** A similar pattern was also observed for the Cookie Theft task. There were no group level differences in total word count (χ^2^ (2, N = 73) = 5.32, p = .070) or content word count (χ^2^ (2, N = 73) = 3.79, p = .150, **Table 3)**. Total word count and content word count correlated between the Famous Face Task and the Cookie Theft Tasks for patients, but not HCs **(Figure S1A and B)** Together, these findings suggest that patients had similar speech output compared to healthy controls on both tasks, and that poorer recall of famous faces seen in LTLE patients could not be explained by decreased overall speech output.

**Table 3.** Famous Face Results (n=73)

|  |  | LTLE (n = 28) | RTLE (n = 18) | Healthy (n = 27) | Total (n = 73) | *P value* |
| --- | --- | --- | --- | --- | --- | --- |
| Famous Face Recognition Accuracy | Mean (%) | 93.04 | 93.08 | 95.80 | 94.09 | 0.78 |
|  | (± SD) | 9.84 | 15.07 | 5.14 | 9.61 |  |
| Speech Duration (sec) | Mean (%) | 644.17 | 986.60 | 737.00 | 766.70 | 0.178 |
|  | (± SD) | 380.1 | 706.42 | 413.01 | 502.05 |  |
| Human-Generated Keywords | Mean | 0.24 | 0.30 | 0.31 | 0.28 | *0.007 |
|  | (± SD) | 0.12 | 0.10 | 0.12 | 0.12 |  |
| Automated-Keywods | Mean | 0.24 | 0.31 | 0.32 | 0.29 | *0.007 |
|  | (± SD) | 0.12 | 0.10 | 0.13 | 0.12 |  |
| Word Count | Mean | 798.36 | 1394.17 | 1137.41 | 1070.67 | 0.37 |
|  | (± SD) | 430.07 | 1292.47 | 900.93 | 901.97 |  |
| Content Words | Mean | 291.36 | 497.94 | 426.78 | 392.38 | 0.34 |
|  | (± SD) | 174.08 | 478.55 | 326.46 | 333.34 |  |
| * = Significant Results |  |  |  |  |  |  |

**Table 4.** Cookie Theft Results (n=53)

|  |  | LTLE (n = 17) | RTLE (n = 13) | Healthy (n = 23) | Total (n = 53) | *P value* |
| --- | --- | --- | --- | --- | --- | --- |
| Speech Duration (sec) | Mean | 80.25 | 84.80 | 91.68 | 86.33 | 0.80 |
|  | (± SD) | 59.71 | 45.57 | 56.54 | 54.35 |  |
| Word Count | Mean | 128.59 | 153.54 | 186.83 | 159.98 | 0.20 |
|  | (± SD) | 99.14 | 108.29 | 109.45 | 107.00 |  |
| Content Words | Mean | 59.12 | 66.77 | 80.35 | 70.23 | 0.16 |
|  | (± SD) | 45.80 | 43.06 | 47.50 | 46.00 |  |

### Famous Face recall performance correlated with MOCA and RVLT scores for TLE subjects

There was a positive significant relationship between FF recall performance and MOCA scores as scored by the human-generated (ρ= .327, p= .029) and automatically-generated dictionaries (ρ= .422, p= .004) for TLE subjects, but not HCs **(Fig 4A, B)**. For TLE subjects with neuropsychological testing (n=18), there was a positive, statistically significant relationship between RVLT score and FF recall performance as scored by the human generated (r=0.501, p=0.018) and automatically-generated dictionary (ρ= .538, p= .001) **(Fig 4C, D)**.

**Figure 4.**
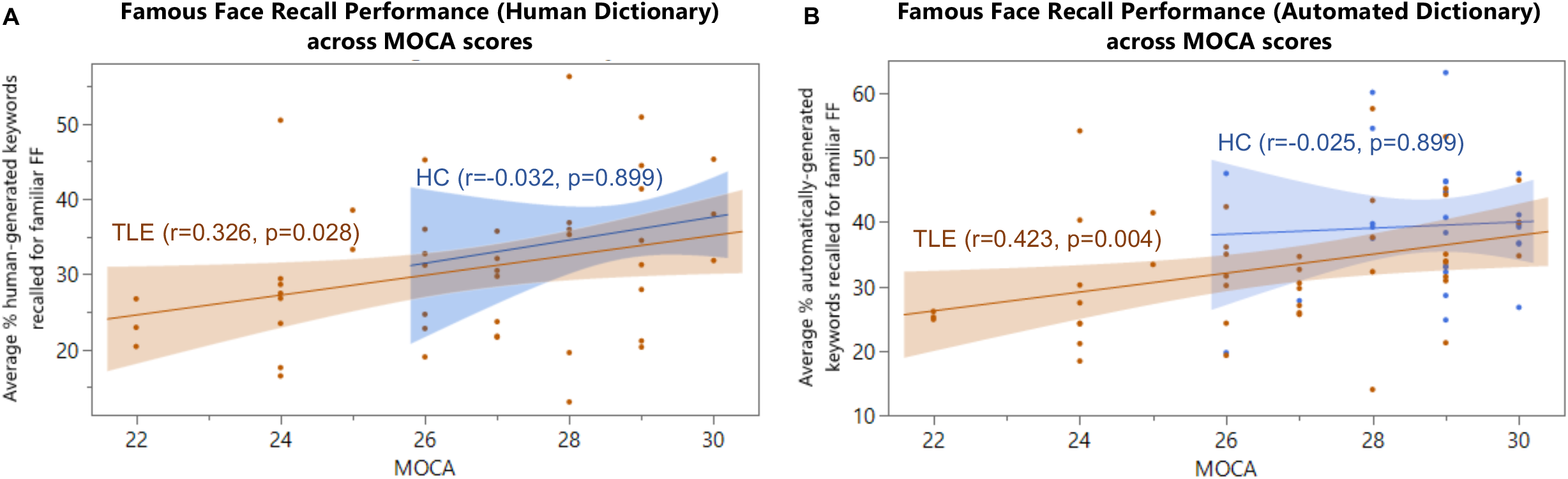
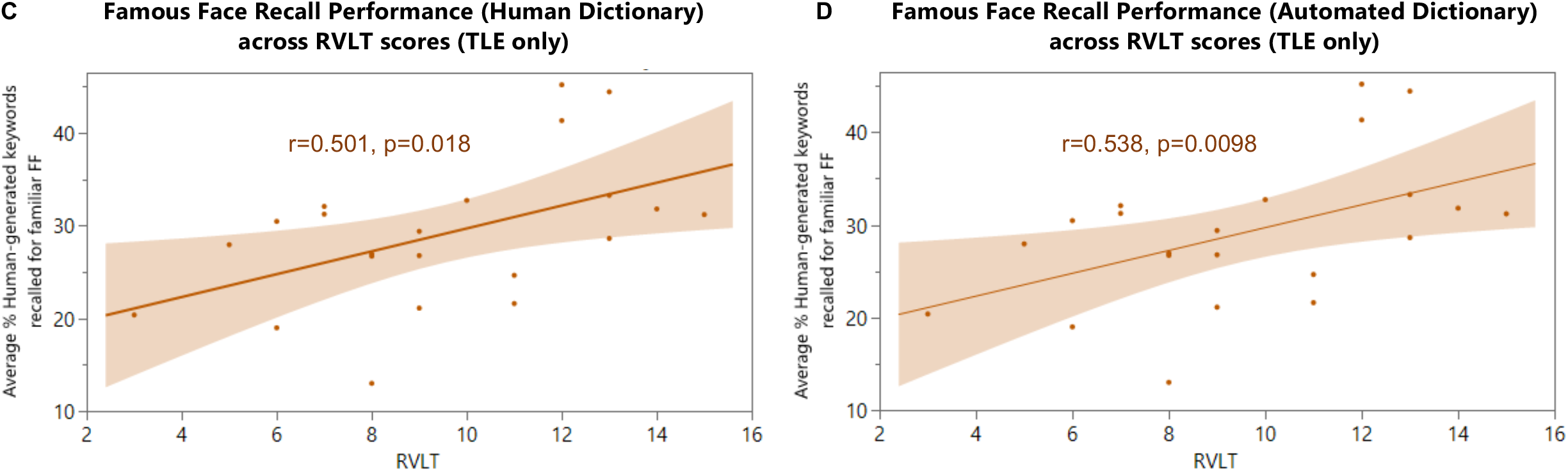
Famous Face Recall Performance correlates with MOCA scores for TLE patients (N=45), but not HCs (N=27)

## DISCUSSION

In summary, patients with left temporal lobe epilepsy generated fewer biographical details of celebrity faces compared to healthy controls or right temporal lobe epilepsy patients, as measured by human and automated analysis of spontaneous spoken recall. Poorer memory recall was not merely an artifact of decreased speech output in the LTLE group, as there were no group differences in speech duration, total word count or content word count during FF recall or the control CTT task.

Our novel approach replicates previous Famous Face recall findings^20,21,26^ and extends them by demonstrating how automated approaches applied to naturalistic behavior can generate a meaningful and quantifiable cognitive measurement. We illustrate how complex human behavior can be scored in a precise, quantitative, and efficient manner. Additionally, we demonstrate how memory can be disambiguated from language. Importantly, FF memory scores derived from spontaneous recall correlate with standardize cognitive and memory tests (i.e., MOCA and RVLT) scores, but display a much wider range of memory performance, and therefore could measure more subtle memory decline. Indeed, cognitive heterogeneity has been well-described in the epilepsy neuropsychological literature as demonstrated in recent studies confirming the presence of multiple cognitive phenotypes in patients with TLE ^34^.

We envision that these methods could eventually be applied to other patient populations at risk for memory decline. Recording and analyzing samples of patient speech during the clinical interview could provide a snapshot of memory and language performance. NLP metrics applied to spoken recall, and extemporaneous speech, could complement existing neuropsychological methods that provide normative data. Furthermore, these methods could provide serial measurements of memory and language with less concern of practice effect.

Prior machine learning methods have been applied to patients with psychiatric disorders and probable Alzheimer’s disease. Acoustic, lexical, and syntactic features can distinguish patients from healthy controls. In psychiatry, patients with PTSD can be distinguished from HCs from acoustic features of speech (e.g., monotony) with high accuracy ^35^. Linguistic features of speech, including semantic density and talk about voices and sounds can predict conversion to psychosis in a high-risk youth cohort with >90% accuracy ^36^. Lexical features such as word repetition, revisions, filler words, utterances, word replacement, and phonemic paraphasias distinguish AD speech from healthy speech ^15,37^. Automatic speech analysis has been applied to identify subtypes of AD, such as primary progressive aphasia ^38^. While these studies show the enormous potential of NLP to extract speech-based features to aid neuropsychiatric diagnosis, we are unaware of any studies that have demonstrated how to assess accuracy and depth of memory through a top-down (human-generated) and bottom-up (automatically-generated, text-driven from healthy subjects) method.

Moreover, to our knowledge, ours is the first application of NLP methods to study speech in epilepsy patients and demonstrates how speech output can be disambiguated from verbal recall. Prior work in epilepsy has focused on extracting textual information from the electronic medical record (EMR). These analyses have demonstrated high accuracy to classify non-epileptic events vs. seizures, presence, or absence of epilepsy, focal versus generalized epilepsy, surgical candidacy, or presence or absence of risk for Sudden Death in Epilepsy (SUDEP) risk ^39–44^.

### Limitations

Limitations of our study include demographic differences between our HC control group and our LTLE patients. HC patients were younger than LTLE patients and had higher MOCA scores (by eligibility criteria). However, we do not think that LTLE memory differences are due primarily to these differences, as the RTLE group which was matched to LTLE group in age and MOCA score also demonstrated superior remote memory. We also acknowledge limitations generalizing the Famous Faces task for clinical purposes. We found that recognition of the twenty celebrity faces was near ceiling for all groups, suggesting a high degree of exposure. Yet, several of the celebrities who were considered prominent in the decade prior to task inception (2018) were not recognizable by the majority of participants. These results suggest a very high degree of exposure to celebrity personalities, that can shift quickly over the span of years. Future adaptations of task stimuli could start with description of a commonly experienced event, such as a film or a news summary, then test for recall after serial delays. Approaches utilizing high performing language and AI models that assimilate the vast amount of information into cohort-specific test stimuli are another possibility.

### Future Directions and Summary

The application of NLP to cognitive testing in epilepsy mirrors the shift in cognitive neuroscience to embrace more naturalistic memory paradigms. Task stimuli are moving away from presentation of words and objects to richer, continuous experiences such as film watching ^10,45^, story listening ^8^, and physical exploration ^11^. The study of complex behavior requires computational analysis to efficiently distill large amounts of data into interpretable and quantifiable measurements. With the rise of artificial intelligence, the detection of subtle memory impairments that may be invisible to conventional testing is possible.

Future work can employ more sophisticated models of language analysis, such as BERT, that have been pre-trained on large datasets of text gleaned from the internet. Larger sample sizes of healthy controls are required to create more robust automated data dictionaries. Additionally, a more detailed analysis of chronological or semantic features of memory could be possible. Finally, to grade memory accurately and on a larger scale, testing would require comparison to a verifiable data source. While famous faces, historical events, and media events are publicly experienced events that can be verified, but their recall is expected to be highly subject to the cultural and educational background of the subject. The accuracy of the patient medical interview could be confirmed by a family member and scored by the number of details remembered.

In summary, NLP methods can be applied to study complex behavior in humans, as in spontaneous recall of famous faces. NLP approaches could be applied in an efficient manner to detect cognitive impairment at the earliest, actionable stage in patients with temporal lobe epilepsy and subjective cognitive impairment.

## Acknowledgements

We thank Ayelet Rosenberg MS for editing and referencing and Kristie Bauman MD, Josh Larocque MD PhD, Hunaid Hassan MD, and Amadou Camara MD for assisting with patient screening.

## Funding

NIH R01NS127954 (AL)

NIH K23NS104252 (AL)

NYU FACES (AL)

