## Supplemental Figures for "Natural Language Processing Applied to Spontaneous Recall of Famous Faces Reveals Memory Dysfunction in Temporal Lobe Epilepsy Patients"

### Table S1. Human and Automated Keyword Dictionaries

**Supplementary Table 1.** Human-generated keywords

| Famous Face | George Cloone | Lisa Kudrow | Prince | Britney Spears | Sarah Palin | Michael Phelps | Nicolas Cage | Demi Moore | Bill Clinton | Mike Tyson | Lucy Liu | Angelina Jolie | Madonna | Eddie Murphy | Cameron Diaz | Tiger Woods | Lady Gaga | Robin Williams | Nicole Kidman | George W. Bush |
|---|---|---|---|---|---|---|---|---|---|---|---|---|---|---|---|---|---|---|---|---|
| | George Clooney | Lisa Kudrow | Prince | Britney Spears | Sarah Palin | Michael Phelps | Nicolas Cage | Demi Moore | Bill Clinton | Mike Tyson | Lucy Liu | Angelina Jolie | Madonna | Eddie Murphy | Cameron Diaz | Tiger Woods | Lady Gaga | Robin Williams | Nicole Kidman | George W. Bush |
| | actor | actor | singer | singer | Alaska | swimmer | actor | actress | president | wrestler | actress | actress | singer | actor | Actress | pro | singer | comedian | actress | president |
| | married | actress | guitar | shave | Tina Fey | Olympic | movie | bruce willis | united states | ear | TV | Brad Pitt | song | comedian | movie | golfer | actress | actor | tv | US |
| | Brad | Phoebe | musician | father | SNL | medal | national treasure | ashton kutcher | hillary clinton | boxer | Charlie's Angels | Children | pop | SNL | charlie's angels | cheat | pop | depression | show | republican |
| | Oceans | Friends | drug | Mickey Mouse | politics | gold | action | married | marries | bitten | movie | Adopt | children | saturday night live | | African-American | pop artist | doubtfire | movie | Barack Obama |
| | movie | TV | pass | Vegas | governor | endorse | face off | singer | chelsea | lisp | | married | adopted | funny | | wife | fashion | mrs. doubtfire | keith | Nine-eleven |
| | Amal | Hollywood | range | Toxic | Vice President | U.S. | hollywood | daughter | affair with monica | tattoo | | divorce | like a virgin | movie | | dirnk | artist | suicide | keith urban | barbara |
| | American | sitcom | Purple Rain | Hit Me | McCain | record | steals the constitution | | intern | heavyweight | | movie | material girl | standup | | PGA | music | passed | tom cruise | iraq |
| | bachelor | famous | song | Oops | Republican | tall | | | impeached | match | | Jennifer Aniston | eighties | donkey | | scandal | a star is born | aladdin | australia | afghanistan |
| | kid | funny | movie | music | Russia | body | | | controversy | | | Mr. and Mrs. Smith | dance | shrek | | | bradley cooper | mork and mindy | | war |
| | child | comic | fashion | Justin Timberlake | election | marijuana | | | democrat | | | salt | | coming to america | | | red carpet | | | upside down |
| | baby | | Minneapolis | married | 2012 | | | | Bush | | | spy | | | | | | | | throw a shoe |
| | famous | | Minnesota | two-thousands | lipstick | | | | scandal | | | | | | | | | | | |
| | attractive | | falsetto | mental | Bulldog | | | | | | | | | | | | | | | |
| | Human Rights | | vocal | Kevin Federline | daughter | | | | | | | | | | | | | | | |
| | lawyer | | The Artist formerly | | controversial | | | | | | | | | | | | | | | |
| | | | Red Corvette | | 2008 | | | | | | | | | | | | | | | |
| | | | Kiss | | accent | | | | | | | | | | | | | | | |

**Supplementary Table 1.** Automatically-generated keywords

| Famous Face | George Cloone | Lisa Kudrow | Prince | Britney Spears | Sarah Palin | Michael Phelps | Nicolas Cage | Demi Moore | Bill Clinton | Mike Tyson | Lucy Liu | Angelina Jolie | Madonna | Eddie Murphy | Cameron Diaz | Tiger Woods | Lady Gaga | Robin Williams | Nicole Kidman | George W. Bush |
|---|---|---|---|---|---|---|---|---|---|---|---|---|---|---|---|---|---|---|---|---|
| | billions | Phoebe Buffay | Corvette | visual | John McCain | legs | Face Off | Bruce Willis | Bush | lisp | wanna say | children | looks | standup comedy | Barrymore | sort | singer | had depression | husband | image |
| | space | Phoebe on Friend | drug overdose | mental | lipstick | body | see much | daughters | office | speaks | agent | Jennifer Aniston | Virgin | laughs | spice girls | car | award | voices | Keith Urban | shoe |
| | famous actor | TV show called Fri | guitar player | trying | running mate | won | famous movies | Friends | Democratic | heavyweight | actress | Mr. and Mrs. Smith | Material Girl | funny | coming | accident | fashion | comedy | actress | George W. Bush |
| | Brad Pitt | sitcom | very high | father | Tina Fey | Olympian | hidden | Demi Moore | impeached | ear | Charlies Angels | lips | Janet Jackson | actor | actress | scandals | Star Is Born with B | actors | Tom Cruise | United States |
| | movies | other guys | falsetto | Madonna | ran | gold medals | point break | Courtney Cox | United States | face tattoo | Lucy Liu | actress | famous singer | Coming | Charlies Angels | PGA | meat dress | Mork | Nicole Kidman | father |
| | married | Lisa Kudrow | mansion | Hit Me Baby | Republican | Olympics | national | | president | boxing | | adopted | charity | Saturday Night Liv | Cameron Diaz | cheated | stage name | Mindy | | Barack Obama |
| | lawyer | | purple | More Time | Vice President | swimming | action movies | | President | boxer | | Jessica Alba | seventies | SNL | | golf player | bars | Aladdin | | Republican |
| | Oceans | | people liken | stars | Sarah Palin | swimmer | actor | | Monica Lewinsky | Boxer | | Brad Pitt | much music | comedian | | tournaments | went | one knew | | served |
| | Amal | | sang | Vegas | Alaska | Michael Phelps | National Treasure | | Hillary Clinton | Mike Tyson | | Angelina Jolie | halftime | Eddie Murphy | | wife | music | passed | | fourty |
| | George Clooney | | songs | singer | | | Nicolas Cage | | Bill Clinton | | | | songs | | | won | A Star Is Born | sad | | third |
| | | | Minnesota | head | | | | | | | | | dance | | | golfer | Bradley Cooper | Mrs. Doubtfire | | Nine Eleven |
| | | | musician | songs | | | | | | | | | pop | | | Tiger Woods | real name | committed suicide | | President |
| | | | passed | Mickey Mouse Club | | | | | | | | | eighties | | | | songs | comedian | | Iraq |
| | | | Red | conservatorship | | | | | | | | | Madonna | | | | pop | Robin Williams | | war |
| | | | Purple Rain | shaved | | | | | | | | | | | | | Lady Gaga | | | George Bush |
| | | | Prince | pop | | | | | | | | | | | | | | | | president |
| | | | | Britney Spears | | | | | | | | | | | | | | | | |

### Figure S1A. FF Word Count correlates with Cookie Theft Word Count for TLE Patients, but not for Healthy Controls

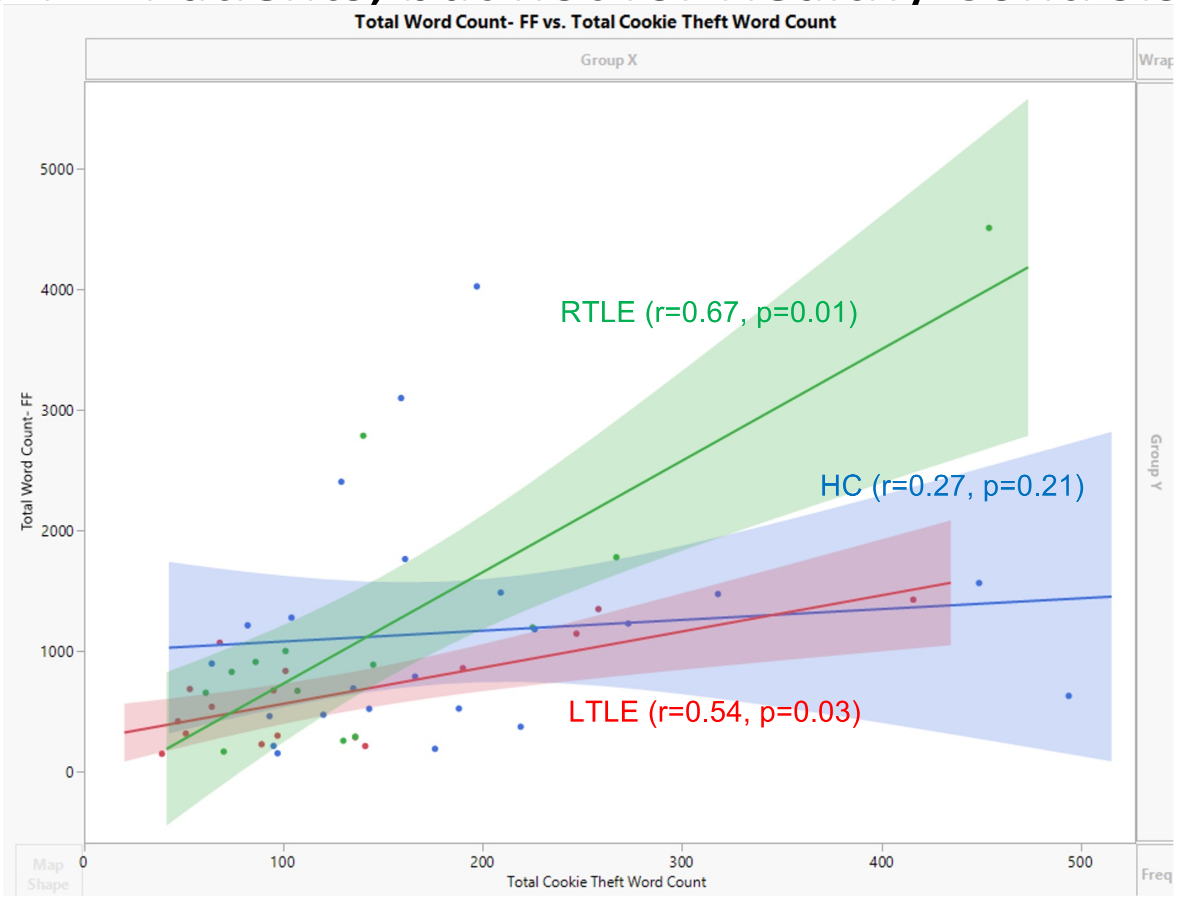

Figure S1B. FF Content Word Count correlates with Cookie Theft Content Word Count for LTLE only, not for HC or RTLE

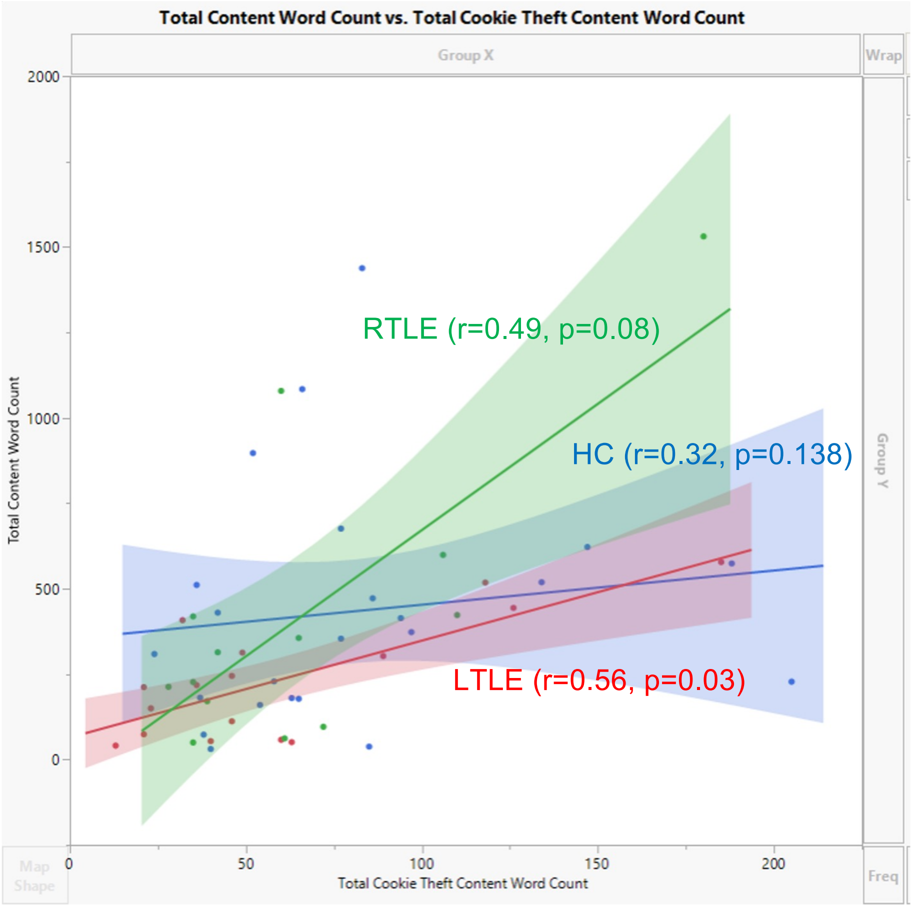

#### Figure S2: FF Recognition in a Multiple Choice Task between Groups

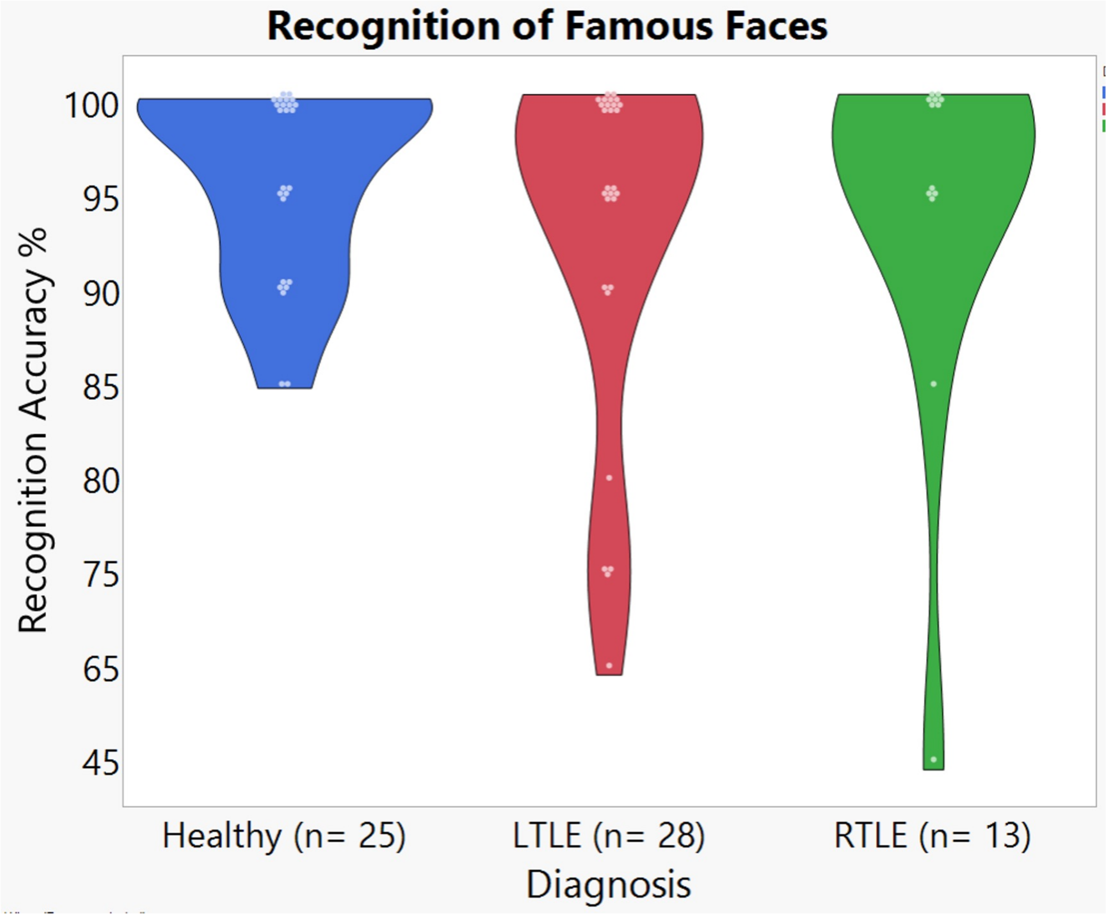

### Figure S3. Famous Face Recall as Scored by a Human and Automated Dictionary correlated with MOCA Scores, by group.

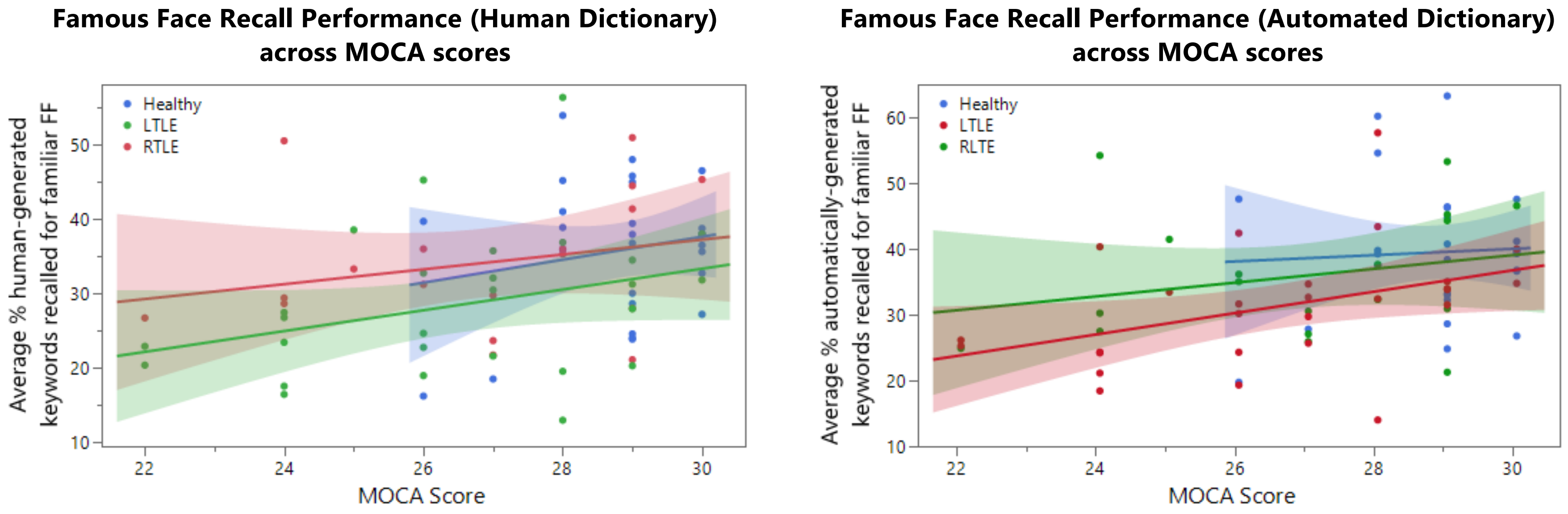
